# A large self-transmissible plasmid from Nigeria confers resistance to multiple antibacterials without a carrying cost

**DOI:** 10.1101/545939

**Authors:** Rubén Monárrez, Molly Braun, Olivia Coburn-Flynn, João Botelho, Babatunde W. Odeotyin, Jose I. Otero-Vera, Naa Kwarley Quartey, Luísa Peixe, A. Oladipo Aboderin, Iruka N. Okeke

**Author notes:** Correspondence to: Iruka N Okeke, Department of Pharmaceutical Microbiology, Faculty of Pharmacy, University of Ibadan, Nigeria.

## Abstract

Antimicrobial resistance is rapidly expanding, in a large part due to mobile genetic elements. We screened 94 fecal fluoroquinolone-resistant *Escherichia coli* isolates from Nigeria for six plasmid-mediated quinolone resistance (PMQR) genes. Sixteen isolates harbored at least one of the PMQR genes and four were positive for *aac-6-Ib-cr*. In one strain, *aac-6-Ib-cr* was mapped to a 125 Kb self-transmissible IncFII plasmid, pMB2, which also bears *bla*_*CTX-M-15*_, seven other functional resistance genes and multiple resistance pseudogenes. We hypothesized that pMB2 had been selected by antimicrobials and that its large size would confer a growth disadvantage. However, laboratory strains carrying pMB2 grew at least as fast as isogenic strains lacking the plasmid in both rich and minimal media. We excised a 32 Kb fragment containing the *sitABCD* and another putative transporter, *pefB*, a *papB* homolog, and several open-reading frames of unknown function. The resulting 93 Kb mini-plasmid conferred slower growth rates and lower fitness than wildtype pMB2. Trans-complementing the deletion with the cloned *sitABCD* genes confirmed that they accounted for the growth advantage conferred by pMB2 in iron-depleted media. The mini-plasmid additionally conferred autoaggregation and was less transmissible and both phenotypes could be complemented with a *pefB* clone. pMB2 is a large plasmid with a flexible resistance region that contains multiple loci that can account for evolutionary success in the absence of antimicrobials. Ancillary functions conferred by resistance plasmids can mediate their retention and transmissibility, worsening the trajectory for antimicrobial resistance and potentially circumventing efforts to contain resistance through restricted use.

## Introduction

Ciprofloxacin was introduced into clinical medicine in the 1980s. It however only became affordable, and therefore widely available in Nigeria in the 2000s, once its patent expired. Fluoroquinolone resistance subsequently emerged in Nigeria (1) and poses an enormous clinical challenge because ciprofloxacin and other orally active quinolones represent the last accessible therapeutic option for many infected patients. High-level fluoroquinolone resistance is most commonly attributable to multiple mutations in the quinolone-resistance-determining regions of quinolone targets *gyrA* and *parC*. In non-mutator *E. coli* strains, the mutation rate is 10^-9^ or lower so that step-wise mutation to resistance is predicted to be slow. Resistance evolved quickly in Nigeria, as in many other parts of the world, supporting the idea that low-level resistance mechanisms may help to protect susceptible strains until they accumulate the 2-4 mutations required for high-level resistance (2). We have previously shown that early quinolone resistant isolates showed elevated efflux capacities (1) but the contribution of plasmid-mediated quinolone resistance (PMQR) in Nigeria and other African countries has been investigated only minimally.

Plasmid-encoded resistance genes most significant in quinolone resistance can be grouped into three functional categories. The *qnr* genes encode members of a pentapeptide-repeat family and confer resistance through direct binding of their gene products to the quinolone targets and thus preventing interactions between the drug and the target enzyme (3, 4). *qnr* genes are suspected to be the most common PMQR genes globally (4). A second category of quinolone-resistance genes, *qepA* and *oqxA,* encode efflux pumps. QepA is a member of the 14-transmembrane-segment major facilitator family of transporters and functions as a proton antiporter efflux pump system that is especially efficient at exporting quinolones but which can also export other antimicrobials including erythromycin, acriflavine, and ethidium bromide. The plasmid-encoded efflux pump, encoded by the *oqxAB* gene is a member of the resistance-nodulation-division family of multidrug efflux pumps and was first identified as conferring resistance to the porcine growth enhancer quinoxaline-di-N-oxide, a compound that also inhibits DNA synthesis (5). *oqxAB,* can also confer resistance to chloramphenicol in addition to the quinolones nalidixic acid and ciprofloxacin (5, 6). The *oqxA* gene appears to be rare among isolates identified in Denmark, Sweden, and South Korea. Recent reports suggest that it is common in South Africa and potentially other parts of the continent but there are few studies performed in sub-Saharan Africa have sought these genes (5, 7-11).

The third and final PMQR category contains only one member. *aac(6’)-Ib-cr* is an aminoglycoside acetyltransferase (12). The standard *aac(6’)-Ib* allele confers aminoglycoside resistance but two SNPs generate a *aac(6’)-Ib-cr* allele that confers resistance to ciprofloxacin and norfloxacin, which have unsubstituted piperazinyl groups, but not to other quinolones (12). The *aac(6’)-Ib-cr* allele was initially identified by Robicsek et al. from clinical *E. coli* isolates originating in China (4). Like the other PMQR genes, with the exception of *oqxA*, *aac(6’)-Ib-cr* has previously been identified in isolates from West Africa (13). On its own, *aac(6’)-Ib-cr* confers only low-levels of resistance to ciprofloxacin but the gene enables target-site mutants to survive at drug concentrations achieved in the clinic. Additionally, *aac(6’)-Ib-cr* is frequently found on plasmids with other resistance genes including *qnr* genes, and *qepA*, as well as ß-lactamases (2, 4).

Although the aforementioned genes have been found throughout the globe there remains a dearth of information on the nature of quinolone resistance in Africa and the relative importance of PMQR genes on the continent. We recently examined antimicrobial resistance in *Escherichia coli* from mother-infant pairs in Nigeria and found that 94 of 1,098 (8.6%) of the isolates were fluoroquinolone resistant (14, 15). We sought to determine the proportion of these strains carrying PMQR genes and the identity of these genes. While we have previously described a plasmid carrying *qnrS1* from a Nigerian isolate (16), no local information is available about the context of *aac(6’)-Ib-cr*. We therefore sequenced and characterized a *aac(6’)-Ib-cr-*bearing plasmid identified in the course of the study.

## Materials and methods

### Strains

Strains used in this study are listed in Table 1. Commensal *E. coli* were collected from Nigerian mother-infant pairs where the infant suffered from diarrhea as previously described (14). Altogether stool samples from 134 mothers-infant pairs, yielded 1,098 *E. coli* isolates for evaluation. Routine culture of bacteria was performed aerobically at 37 ^°^ C in Luria broth supplemented with chloramphenicol (30 μg/mL), ampicillin (100 μg/mL), tetracycline (25 μg/mL), or neomycin (50 μg/mL) where applicable. Strains were maintained at −70°C in Luria broth:glycerol 1:1.

**Table 1.** Strains used in this study.

|  | Genotype and description | Reference or Source |
| --- | --- | --- |
| M63c | *E. coli* wt, S<sup>R</sup>, Nal<sup>R</sup>, Sul<sup>R</sup>, Amp<sup>R</sup>, Cip<sup>R</sup>, Te<sup>R</sup>, W<sup>R</sup>, K<sup>R</sup> | (14) |
| M63a, M63b, M63d, M63d | *E. coli* isolates recovered from the same specimen as M63c | (14) |
| C63a, C63c | *E. coli* isolate obtained from a specimen collected on the same day, from the infant of research participant whose stool yielded M63c | (14) |
| DH5α | F<sup>-</sup> ø80dlacZΔM15 Δ(lacZYA-argF)U169 *deoR recA1 endA1 hsdR17(rK<sup>-</sup> mK<sup>+</sup>) phoA supE44 λ– thi-1 gyrA96 relA1* | Invitrogen |
| C600 Nal<sup>R</sup> | Nalidixic acid-resistant derivative of C600. | (69) |
| NCTC 10418 | Susceptibility testing control |  |
| ATCC 35218 | Susceptibility testing control |  |
| ORN172 | *thr-l leuB thi-1 A(argF-lac)U169 maL1l xyl-7 ara-13 26 mtl-2 gal-6 rpsL tonA2 supE44 Δ(fimBEACDFGH);* | (66) |
| EC1502 | Rifampicin-resistant, plasmid free *E. coli* strain | University of Bradford |
| HS | Commensal *E. coli* strain that has been shown to colonize non- | (70) |
|  | virulently in human volunteer studies |  |

### Antimicrobial Susceptibility Testing

Antimicrobial susceptibility testing was performed using the Clinical and Laboratory Standards Institute (CLSI) disc diffusion method and interpreting zone diameters in accordance with CLSI guidelines in WHONET software version 5.3 (17). Discs used contained ampicillin (10µg/ml), streptomycin (10µg), trimethoprim (5µg), tetracycline (30µg), nalidixic acid (30µg), chloramphenicol (300 µg), sulphonamide (1000 µg) and ciprofloxacin (5µg) (Oxoid/ Remel). *E. coli* ATCC 35218 and DH5αE (Invitrogen) were used as control strains. Minimum inhibitory concentrations (MICs) for nalidixic acid and ciprofloxacin were determined using E-test (bioMérieux) on Mueller-Hinton (MH) agar. Kanamycin MICs were determined by agar dilution on Mueller-Hinton Agar.

### General microbiology and molecular biology procedures

The Promega Wizard® kit was used for genomic DNA extractions and plasmids under 20 Kb were extracted using a Qiagen® MiniPrep Kit. Naturally-occurring low copy number larger plasmids were extracted, after growth of host bacterium in Terrific Broth and induction with chloramphenicol, by a modified boiling protocol for small scale preparations (18, 19) and using the Qiagen® Large Construct Kit for large-scale preparations. Table 2 lists the plasmids used in the study, their relevant properties, and their sizes. Plasmids were electroporated into *E. coli* host strains using a Bio-Rad micropulser according to manufacturers’ instructions. Sequences of oligonucleotide primers used in the study are listed in Supplementary Table 1. Amplification cycles began with a two-minute hot start at 94°C followed by 30 cycles of denaturing at 94°C for 30 s, annealing at 5°C below the melting temperature (unless otherwise specified) for 30 s, and extending at 72 °C for one minute per kilobase of DNA to be amplified. *Bst*CI restriction analysis of *aac(6’)-Ib* PCR amplicons was used to determine whether they represented the *aac(6’)-Ib-cr* allele (20). Up to two amplicons from each unique profile were sequenced to confirm PCR-RFLP-based classification. Where necessary, PCR amplicons were TA-cloned into the pGEMT vector (Promega) according to manufacturer’s directions and plasmids were transformed into chemically competent *E. coli* K-12 TOP10 cells for sequencing. Other molecular biology operations were performed using standard procedures (21).

**Table 2:** Plasmids used in this study.

| Plasmid | Description | Size | Reference or source |
| --- | --- | --- | --- |
| pMB2 | Naturally occurring *aac(6')-Ib-cr*-bearing plasmid | 125 Kb | This study and (16) |
| pMB80-2 | Naturally occurring conjugative plasmid from enteropathogenic *E. coli* strain | >100 Kb | (19) |
| pRMKO | Mini-plasmid constructed by deleting a 32,331 bp *Not1* - *Xba1* fragment from pMB2 | 93 Kb | This study |
| pRMC | pBluescript II SK containing a 32,331 bp *Not1* - *Xba1* fragment from pMB2 cloned | 35 Kb | This study |
| pBAD/Thio-TOPO | Arabinose inducible expression vector | 4,454 bp | Invitrogen |
| pBluescript II SK + | High copy number cloning vector | 2,961 bp | Agilent |
| pINKpefB | pefB gene from pMB2 cloned into pBAD/Thio-TOPO | 4,724 bp | This study |
| pSIT1 | Clone of the *Shigella sitABCD* genes |  | (71) |
| pINK2301 | *Shigella sitABCD* genes subcloned from pSIT1 into | 7.1 Kb | This |
|  | pACYC177 |  | study(16) |
| pLMJ50 | Cm<sup>R</sup>; large aggregative adherence plasmid from EAEC strain 60A in which the *impB* gene is replaced with a *cat* cassette | 90 Kb | (60) |
| pGEM-T | Amp<sup>R</sup>; TA-cloning vector | 3,000 bp | Promega |
| pBR322 | Amp<sup>R</sup>, Tc<sup>R</sup> cloning vector | 4,363 bp | NEB |
| pACYC177 | Amp<sup>R</sup>, Km<sup>R</sup> cloning vector | 3,941 bp | NEB |

### Bacterial identification

Bacterial were identified to the species level as described previously (14) and further biotyped on the API20E system (bioMérieux). Genetic identification was performed by 16S ribosomal subunit sequencing, amplifying the gene encoding M63c’s 16S rRNA with primers 10 and 1507R (Supplemental Table 1) in a thermocycler using the cycle at 94°C for 2 m, [94°C for 30s, 59°C for 45s, 72°C for 90s] x36, and 72°C for 10m (22). Multilocus sequence typing (MLST) was performed by amplifying the *adk, fumC, gyrB, icd, mdh, purA* and *recA* genes using the primers of Wirth et al as previously described (23, 24). PCR amplicons for sequencing were size-verified by agarose electrophoresis, cloned into pGEMT, and electroporated into DH5αE (Invitrogen) using a Biorad Micropulser. Plasmids were extracted with a Qiagen Plasmid Mini Kit and Sanger sequenced from M13F and M13R priming sites.

### Shot-gun sequencing and sequence analysis

Whole-replicon shotgun library preparation, Sanger sequencing and assembly of plasmid pMB2 was performed by SeqWright DNA Technology Services (Houston, TX). Preliminary sequence analyses and annotation was performed manually by four authors in Artemis (25) with open reading frames initially predicted using Glimmer (26-28) and basing gene annotations on 98% or greater identity at the nucleotide and amino acid levels if BLAST e-value was 0 (29). Plasmid MLST typing was performed *in silico*, using PlasmidFinder and pMLST versions 2.0 (30). Orf identity was determined using BLAST and Pfam (27, 28). Two automated annotations was performed using BASys (26) and Prokka v 1.13.3 (31). Five authors resolved annotation discrepancies among automated and manual annotations. Antimicrobial resistance genes and their genetic context were searched using Galileo AMR (https://galileoamr.arcbio.com/mara/) (32) and ResFinder (29). Direct and inverted repeats were identified by dot-plot analysis of pairwise FASTA alignments made using the BLAST suite. Open reading frame (Orf) and feature plots were prepared using BLAST ring image generator (33).

### *In vitro* conjugation

*In vitro* conjugation experiments were performed by solid mating. Donors and recipients were cultured in LB with appropriate selective antimicrobials. 0.5 mL of donor and recipient culture, grown overnight with selection, was spun down gently and resuspended in 20 μl of LB without antibiotics. The suspension was spotted onto dried LB plates, allowed to dry at room temperature for 15 minutes and then incubated at 37°C for three hours. The mating reaction was resuspended in 1ml of LB with vortexing and placed on ice to terminate conjugation. After mating, serial ten-fold dilutions of each terminated reaction was made in cold phosphate buffered saline and plated onto plates containing tetracycline (or other appropriate antimicrobials for controls) to select for the plasmid and nalidixic acid, resistance to which is conferred chromosomally in the recipient. Transconjugant colonies were counted after overnight incubation at 37°C and verified by plasmid profiling, phenotype on MacConkey and Eosin methylene blue agars, PCR-RFLP for the *fliC* allele (34), and PCR for donor- and recipient-specific markers. Viable counts of donors and recipient were also performed. The number of transconjugant colonies per donor colony-forming units was computed as the plasmid transfer efficiency (35)

### Growth curves

Growth curves were plotted following growth of each strain in microtiter wells containing 160 µl of media in an Infinite^®^ 2000 Pro Series microplate reader preset at 37°C (Tecan). Commercially available media were prepared as recommended by the manufacturers. Iron-depleted media was prepared by adding deferrated ethylene diamino-o-dihydroxyphenyl acetic acid (EDDA) as described previously (36). Test strains were first cultured overnight in 5 mL of LB media containing appropriate antibiotics where required. Overnight cultures were spun down, resuspended in 5 mL of 1X Phosphate Buffered Saline (PBS), and diluted to an optical density of 0.70 at 595 nm. 10 µl of the diluted cultures were then loaded into assigned wells in the microtiter reader. Absorbance was measured at 595 nm every 30 min preceded by 30 sec of orbital shaking for 24hours. The average optical density of 25 reads per well at each time point was used in the analysis and up to six wells were tested per strain.

### Autoaggregation assay

Autoaggregation was quantified in bacterial-settling liquid culture assays as described by Hasman et al (37). Overnight cultures were prepared in test medium containing added selective antibiotics where necessary. For experiments requiring induction, gene expression was induced for 90 minutes before the start of the assay. At the assay start, cultures of each strain were adjusted to the same optical density at 600nm (OD_600_). Eight ml of each adjusted culture was placed into two separate tubes. One tube remained static and the other was lightly vortexed before each optical density measurement. The tubes were incubated without shaking at 37 °C. At designated time points, 0.5 mL was removed from within 2 cm of the surface of the culture, and the OD_600_ was measured, diluting the sample if required. OD_600_ supernatant measurements for different strains were compared using a t-test.

### Plasmid stability

Plasmid stability was assessed by serially passaging *E. coli* strains carrying pMB2 and its derivatives as described by Sandegren et al (38). Triplicate starting cultures were grown overnight at 37°C in 1mL of LB supplemented with ampicillin or tetracycline. Bacterial cells were washed, resuspended in LB without antibiotics and. An aliquot of 100 µL of washed cells was inoculated into 1 mL of LB and incubated overnight at 37°C. 100 µL of this overnight culture was serial passaged to 1mL of LB was performed daily, approximating 10 generations of growth per passage. At select time points, samples were diluted and plated onto plain MacConkey plates and MacConkey plates containing tetracycline.

### *In vitro* competition experiments

We measured relative fitness in traditional competition assays as outlined by Wiser and Lenski (39). Fitness was computed independently in LB broth and DMEM and *E. coli* strains carrying pMB2 and its derivative plasmids were selected using antibiotic markers present on those plasmids.

## Results and Discussion

### Plasmid-encoded quinolone resistance genes in fluoroquinolone resistant isolates from Nigeria

Of 1,098 fecal *E. coli* isolates originally recovered from 134 mother-infant pairs, 94 (8.6%) were ciprofloxacin-resistant and were screened for six PMQR genes. The PCR screen initially revealed that *aac(6’)-Ib* was common among ciprofloxacin resistant isolates with 37 (39.3%) being positive for the allele as shown in Table 3. However, upon *Bst*I restriction analysis only four of *aac(6’)-Ib* genes (10.8%) proved to be the *aac(6’)-Ib-cr* allele (Table 3). *qnrA* and *qnrB* were not detected at all whilst *qnrS1*, which we have previously detected in a different strain set from Nigeria(16), was found in two of the isolates. Overall, *oqxAB* was detected in nine isolates (including two that were positive for *aac(6’)-Ib-cr*) and was therefore the most common PMQR detected but *aac(6’)-Ib-cr* was the most common quinolone-specific mechanism (Table 3). We therefore went on to characterize M63c, one of the strains bearing *aac(6’)-Ib-cr* alleles.

**Table 3:**
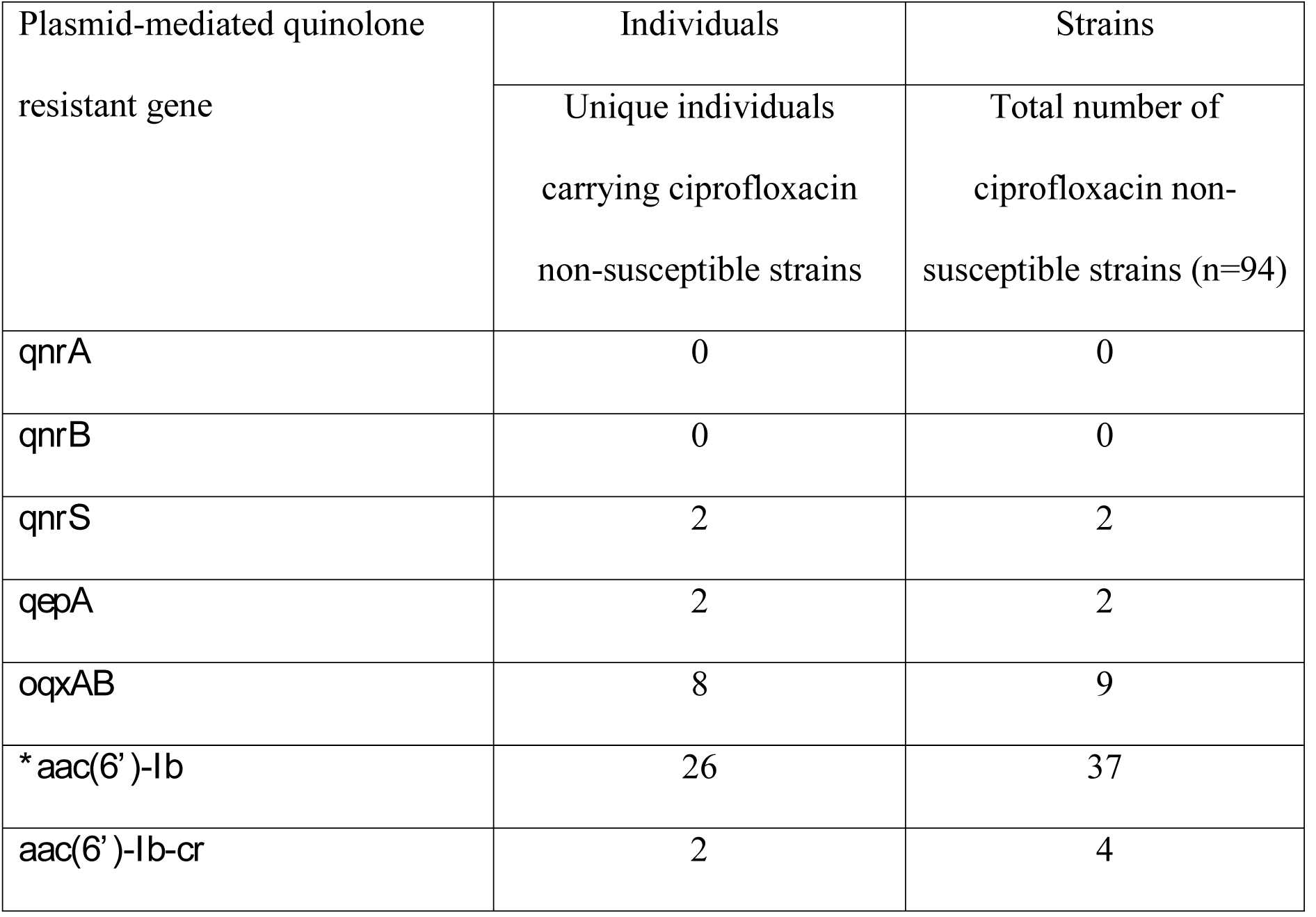
PMQR genes identified by PCR in ciprofloxacin non-susceptible *Escherichia* isolates Includes the PMQR *aac(6’)-Ib-cr* as well as non-ciprofloxacin resistance conferring *aac(6’)-Ib*

Strain M63c was mucoid on MacConkey and eosin methylene blue plates with an API 20E profile pointing to ‘group C *E. coli*’, which are now classified as *Escherichia fergusonnii*. We amplified and sequenced the 16S rRNA gene of M63c and found the sequence (Accession number SUB5013267) was 99% identical to that of *Escherichia fergusonii* type strain ATCC 35469 (Accession number NR_074902.1). However, Maheux et al (40) reported that the *adk, gyrB* and *recA* genes were more useful for speciating *Escherichia* spp. than 16S rRNA. Alleles of those genes carried by M63c were identical to *E. coli* alleles. We additionally sequenced *fumC, icd, mdh* and *purA* genes of the strain and multilocus sequence typed the isolate as an *E. coli* ST167 strain. *E. coli* ST167 strains are encountered around the globe, frequently multiply resistant and often expressing extended-spectrum β-lactamases (ESBLs) (41). We extracted a large plasmid DNA from strain M63c and electroporated it into DH5αE. Transformants were obtained on neomycin (50 mg/ml) plates but not on ciprofloxacin (1 mg/ml). Each transformant showed reduced susceptibility to ciprofloxacin and nalidixic acid as well as resistance to tetracycline, ampicillin, neomycin/ kanamycin, trimethoprim and sulphonamides but not chloramphenicol. All transformants had identical plasmid DNA *Bam*HI, and *Sal*I restriction profiles and susceptibility patterns and carried a single large plasmid, which we termed pMB2.

### Plasmid pMB2

pMB2 was Sanger-sequenced and shot-gun assembled into a 125,782 base pair circular replicon (Genbank Accession number MK370889). pMB2 has a G+C content of 51.73% (Figure 1), which is comparable to that of the *Escherichia coli* chromosome (42) A maximal number of 323 ORFs of approximately 100 or more base pairs in length were predicted but additional evidence supports only 145 of these as encoding real or hypothetical genes. The plasmid contains a full IncFII-type conjugation region of 32 genes including all 25 of the consecutive transfer region genes with known functions in the conjugation process (43).

**Figure 1:**
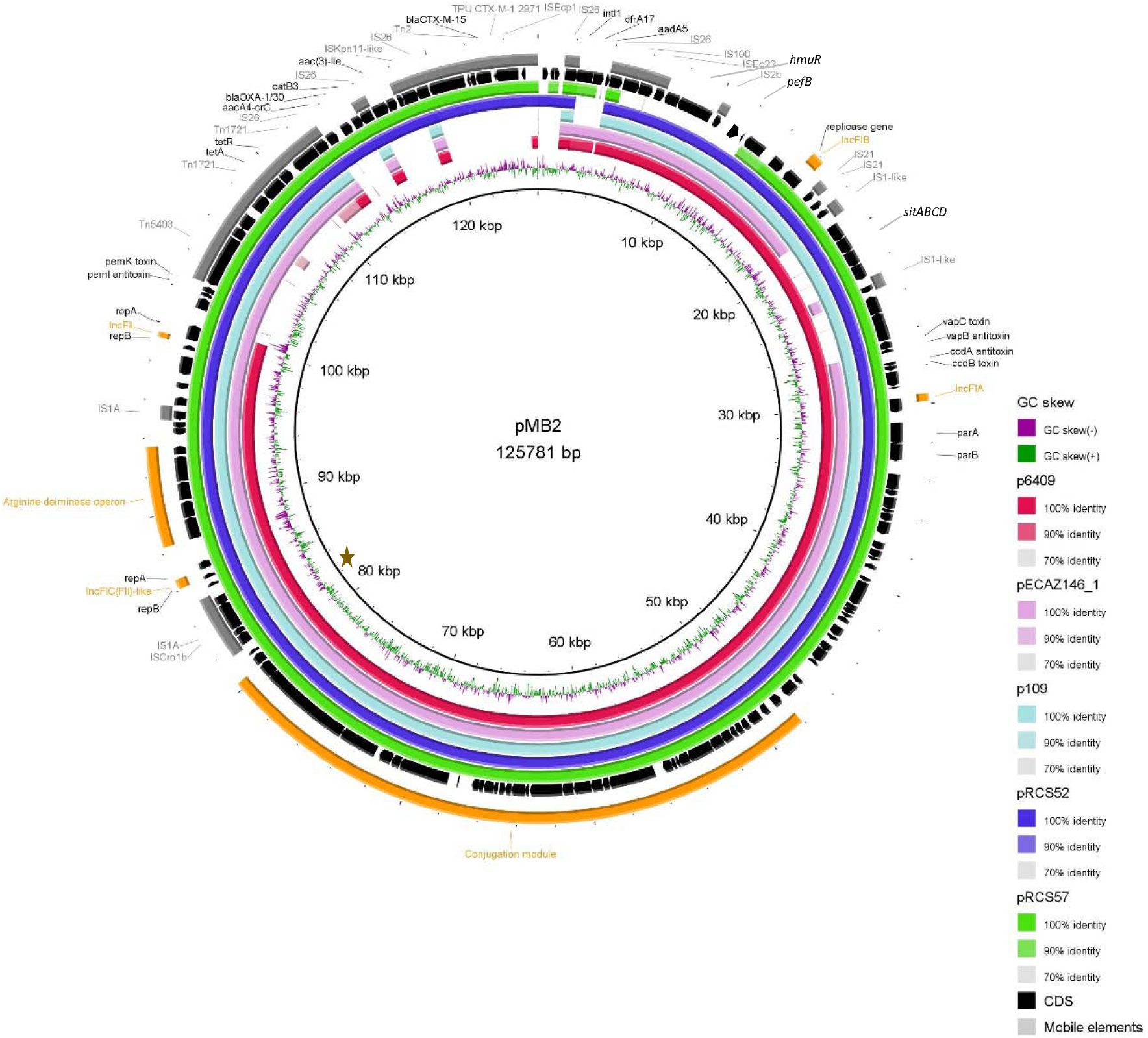
Circular plot illustrating the general features of pMB2. The figure shows the organization of pMB2 orfs in black and a BLASTn comparison between pMB2 and the five best hits *E. coli* plasmids retrieved from NCBI (from the innermost to the outermost circles, plasmids p6409, pECAZ146_1, p109, pRCS52 and pRCS57 with the GenBank accession numbers CP010372.1, CP018990.1, CP023372.1, LO017736.1 and LO017738.1, respectively). A brown star marks the likely location of the origin of replication.

*In silico* replicon typing revealed that the Plasmid Sequence Type of pMB2 is [F36:A4:B1]. The pMB2 plasmid carries replication-associated proteins associated with IncFIA, IncFIB, IncFIC and IncFII plasmids. The IncFIC RepA1 protein is coded by an ORF at 86,659bp on the plasmid, which is close to the major shift in G+C skew seen at 81,000bp and likely marking the origin of replication (Figure 1). pMB2 is rich in mobility genes with at least 26 transposases or transposase-like orfs, as well as up to six other integrases and recombinases encoded. Almost all functional units on the plasmid are flanked by insertion sequences. pMB2 carries genes encoding several toxin-antitoxin systems *ccdA/ccdB*, *vapB/vapC* and *pemI/pemK*. It also carries the SOS response inhibiting proteins PsiA and PsiB. Between 88,324 and 93566 is a complete, six gene arginine deaminase operon, which could potentially catabolize arginine to ammonia.

pMB2 is predicted to encode at least three transport systems. The EamA-like domain domain-containing drug/metabolite and carboxylate/amino acid/amine transporter (Pfam00892; located at 107699-108583), is found in a variety of integral membrane proteins; there is no known function for these genes but they could possibly contribute to antimicrobial resistance. A hypothetical divalent ion transport system (located between 10045 and 7247, on the complementary strand) is composed of two proteins coded in an operon: conserved hypothetical pMB2_00012 a (pfam10670) (44)and HmuR/hemR, which is a Ton-B dependent hemoglobin-hemin receptor localized to the bacteria’s outer membrane (45). The third transport system, and the best studied, is encoded by encoded by *sitABCD* located on the forward strand of the plasmid between 19,195 and 22,644. SitABCD form an effective ferrous iron uptake system earlier described in *Salmonella enterica* and *Shigella* (46, 47). SitA is a periplasmic binding protein, SitB is the ATP-binding component, SitC functions as a permease and SitD is the inner membrane component of the system.

pMB2 bears eight genes conferring resistance to six different drug classes encoded by the plasmid (Figure 2). An IS*26*-composite transposon flanks an incomplete class I integron carrying a *dfrA17* gene (encoding resistance to trimethoprim) and an incomplete *aadA5* gene. The tetracycline resistance gene *tetA* (alongside *tetR*, encoding the repressor of the tetracycline resistance element) are located within the remnants of a Tn*1721* transposon. Next to this region, another IS*26*-composite transposon was identified, carrying an incomplete class I integron with a |*aadA16*|*aac(6’)-Ib-cr*|*bla*_OXA-1/30_|Δ*catB3*| gene cassette array encoding resistance to aminoglycosides, fluoroquinolones, beta-lactams and phenicol. As stated previously, the *aac(6’)- Ib-cr* gene, also named *aacA4-crC,* confers resistance to aminoglycosides and the broad-spectrum antimicrobial ciprofloxacin due to a T to C mutation at nt 283 and a G to T mutation at nt 514. Following *bla*_*OXA-1*_ is *catB3,* which normally inactivates chloramphenicol through acetylation (Shaw, 1983) but which is not active due to the truncated cassette region. Next to the IS*26*-composite transposon is an *aac(3)-II* gene, conferring gentamicin resistance. Finally, a transposition unit carrying *bla*_CTX-M-15_, which confers resistance to extended-spectrum β-lactams, and IS*Ecp1* was found within the remnants of a Tn*2* transposon. This insertion unit is flanked by 5-bp direct repeats (5’-TATGA-3’), suggesting the *en bloc* transposition of these elements within the Tn*2* transposon (Figure 2).

**Figure 2:**
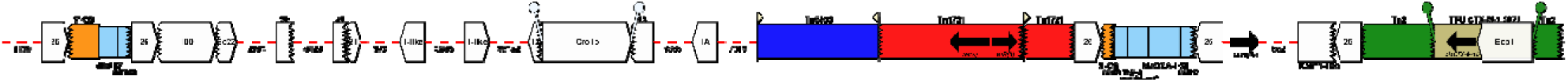
Genetic context of the pMB2 antimicrobial resistance region. Arrows indicate the direction of transcription for genes. Gene cassettes are shown by pale blue boxes, the conserved sequences (5’ and 3’-CS) of integrons as orange boxes and insertion sequences as white block arrows labelled with the IS number/name, with the pointed end indicating the inverted right repeat (IRR). Unit transposons are shown as boxes of different colours and their IRs are shown as flags, with the flat side at the outer boundary of the transposon. Direct repeats are shown as ‘lollipops’ of the same color. The zig-zag line indicates which end of the feature is missing. Gaps >□50□bp are indicated by dashed red lines and the length in bp given.

There have been several reports of co-occurrence of the ESBL gene *bla*_*CTX-M-15*_ and *aac(6’)- Ib-cr* across of the globe, including in Nigeria (48-50). This common co-resistance profile implies that ciprofloxacin use, which is growing (1), also selects for the carriage of ESBL genes and vice versa explaining why IncFII plasmids carrying these genes have traditionally been associated with *E. coli* ST131 and other successful pandemic clones (51, 52). This report appears to be the first one directly confirming a genetic linkage of *bla*_*CTX-M-15*_ and *aac(6’)-Ib-cr* in Nigeria. However the structure of this plasmid in general and the resistance region in particular has been reported from round the world.

pMB2 is similar to many other resistance plasmids in the database that have been isolated from around the world. Figure 1 compares the sequences of four of these plasmids with pMB2 illustrating that the plasmids are almost completely conserved and almost perfectly syntenic. In particular, conjugation and arginine deaminase genes are highly conserved among the plasmids wherease their resistance regions are expectedly the most variable. Only three of the plasmids carry all the putative transport genes, however all of them have at least two of them.

### pMB2 is self-transmissible

While the plasmid as a whole is replete with transposable elements and insertion sequences, and contains multiple antimicrobial resistance genes, some no longer functional, some redundant, the conjugation region of pMB2 is not interrupted. The transfer genes of pMB2 are organized similarly to those of the F plasmid (53) and identical to those on the other five plasmids we compare with pMB2 in figure. Conjugation from DH5α (pMB2) to EC1502 (rifampicin resistant) on solid media occurred at the rate of 1.8 × 10^-5^ to 8 × 10^-6^.

Four other *Escherichia* isolates from the individual from whom strain M63c was isolated had been archived along with two more isolates from her infant (14). As shown in Table 4, strain M63c was the only quinolone-non-susceptible isolate of this set. We screened all six isolates for the presence of the *aac(6’)-Ib-cr* allele and pMB2 and found that they were present in strain M63b, as well as original source strain M63c. While they differed in quinolone resistance, strain M63b and M63c shared otherwise identical susceptibility patterns. We sequenced quinolone resistance determining regions of *gyrA* and *parC* in both strains and found that quinolone-susceptible strain M63b contained no SNPs whilst resistant strain M63c carried Ser83Leu and Asp87Asn mutations in GyrA and Ser80Ile, Glu8Val mutations in ParC. These findings are consistent with reports that demonstrate that those SNPs are sufficient for high-level fluoroquinolone resistance and *aac(6’)-Ib-cr* allele insufficient to confer clinically significant ciprofloxacin resistance but augments resistance conferred by other mechanisms (12, 54). The presence of the plasmid in two different *Escherichia* strains from the same host strongly suggests *in vivo* transmission.

**Table 4:** Antimicrobial susceptibility profiles of Escherichia isolates from the mother-infant pair from whom plasmid pMB2 were recovered and DH5_α_ E with and without pMB2. Data are zone diameters in mm and CLSI interpretations (R= resistant, I = intermediate; S= sensitive) and minimum inhibitory concentrations. A nalidixic acid-resistant derivative of E. coli C600 was used as a quinolone-resistant control.

| | Zone diameters (mm) and CLSI interpretations | | | | | | | | Minimum inhibitory concentrations (μg/mL) | | |
| --- | --- | --- | --- | --- | --- | --- | --- | --- | --- | --- | --- |
|  | C | S | NA | SUL | AMP | CIP | TE | W | NA | CIP | KM |
| DH5αE | 30 S | 23 S | 20 S | 50 S | 40 S | 42 S | 34 S | 22 S | 12 | 0.023 | < 7.5 |
| DH5αE<br>(pMB2) | 30 S | 22 S | 16 I | 44 S | 6 R | 34 S | 9 R | 6 R | 16 | 0.094 | > 30 |
| C600 | 24 S | 17 S | 6 R | 36 S | 23 S | 26 S | 22 S | 30 S | >256 | 0.25 | < 7.5 |
| M63a | 21 S | 14 I | 22 S | 6 R | 6 R | 39 S | 6 R | 6 R | 6 | 0.023 | < 7.5 |
| M63b | 22 S | 13 I | 21 S | 6 R | 6 R | 42 S | 7 R | 6 R | 6 | 0.016 | < 7.5 |
| M63c | 21 S | 10 R | 6 R | 6 R | 6 R | 6 R | 6 R | 6 R | >256 | >32 | > 30 |
| M63e | 21 S | 14 I | 21 S | 6 R | 6 R | 31 S | 6 R | 6 R | 4 | 0.023 | < 7.5 |
| C63a | 21 S | 14 I | 22 S | 24 S | 15 I | 42 S | 17 I | 24 S | 4 | 0.023 | < 7.5 |
| C63c | 22 S | 16 S | 25 S | 30 S | 19 I | 38 S | 21 S | 25 S | 4 | 0.023 | < 7.5 |

### pMB2 does not have a carrying cost

When resistance is conferred by a plasmid of considerable size, the associated metabolic burden on the organism due to the additional energy required in replication and gene expression typically exerts a carrying cost (55). Some research has suggested that this burden is reduced or eliminated through compensatory mutations or acquisitions (56-59). Alternatively, or in addition, plasmid-encoded resistance genes themselves could confer a fitness advantage in the absence of antimicrobials (58). We hypothesized that the large size of pMB2 would slow the growth of laboratory strain DH5α. In rebuttal of our hypothesis, DH5α carrying pMB2 grew at rates that were comparable to or faster than the plasmid-free strain. This occurred in Luria broth (LB) (Figure 3) as well as in nutrient broth, terrific broth, Dubelcco’s Modified Eagle Medium (DMEM, Invitrogen), Davis minimal medium and M9 minimal medium (data not shown). As a control, we used a 92 kb aggregative adherence/*impAB* plasmid, pLMJ50 (60), which did demonstrate a significant carrying cost in DH5α (Figure 3).

**Figure 3:**
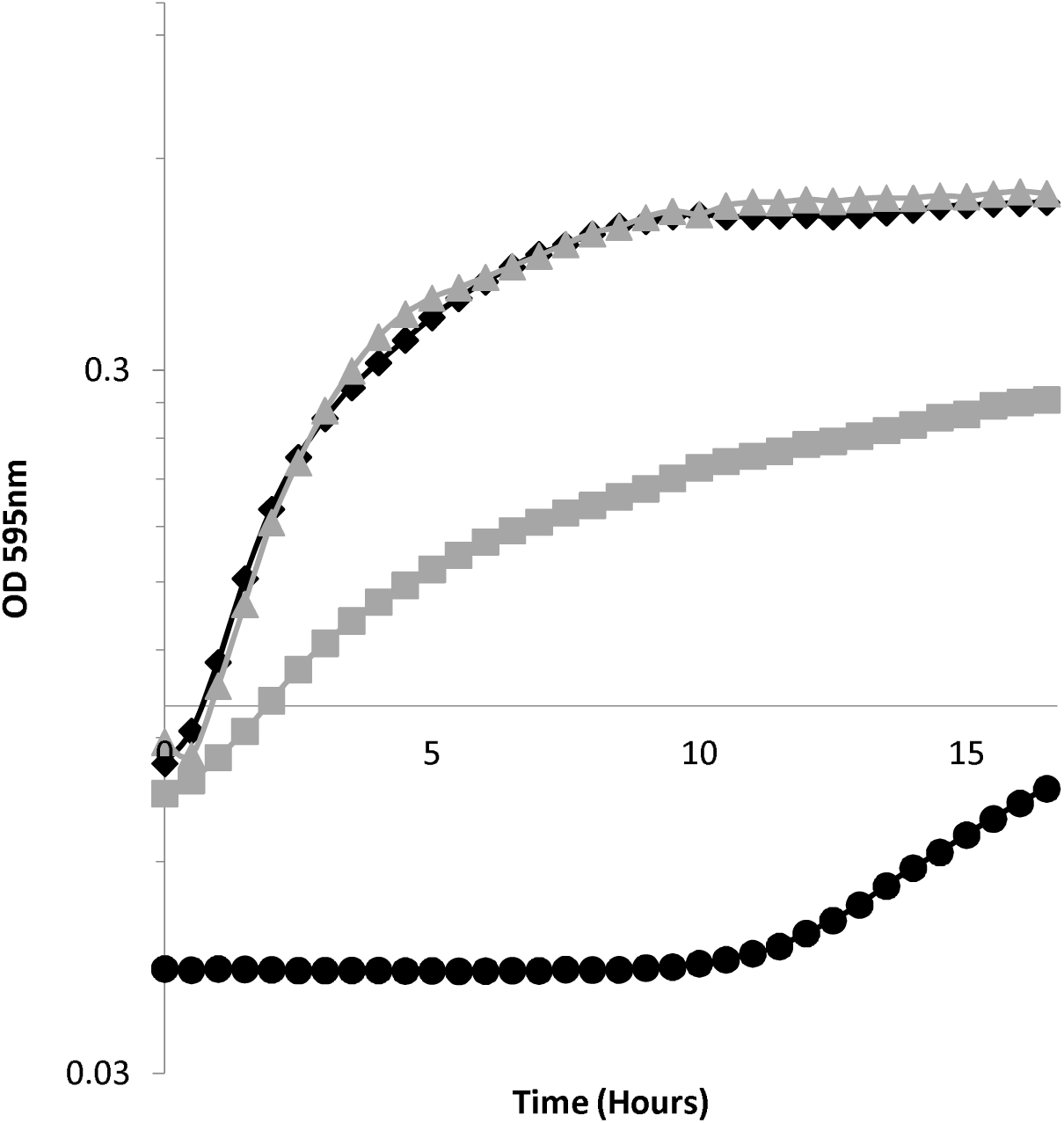
18 h growth time-course of DH5α (black diamonds), DH5α carrying 125 Kb plasmid pMB2 (grey triangles) and 90 Kb plasmid pLMJ50 (grey squares). The black circles represent a no bacteria control.

### Mutational analyses of pMB2

We identified the transport systems operons pMB2_00012/*hemR* and *sitABCD* located at 7,247-8,038 bp and 19,195-22,644 bp respectively on pMB2 as potential candidates for conferring a growth advantage. To determine whether the region harboring these genes offsets a carrying cost of pMB2, a 32,331 bp segment that included these genes was deleted from the plasmid by excision with *Not*1 (NEB), which cuts pMB2 at 122,777 bp and *Xba*1 (NEB), which cuts pMB2 at 29,449 bp. The restricted ends were filled in using *E. coli* DNA polymerase I (NEB) and then ligated on one other to generate a 93,450 bp autonomously-replicating mini-plasmid, pRMKO. The excised 32,331 bp fragment containing the transport system region was cloned into the *Xba*I and *Not*I sites of pBluescript II SK (Agilent) and named pRMC.

Normally, a large deletion in a plasmid should ameliorate carrying costs by reducing the amount of DNA that is replicated and transcription of genes present on the deleted region (55). Therefore, based on size alone, the 93 Kb mini-plasmid, pRMKO, might be expected to confer faster growth than the 125 Kb pMB2 in DH5α but this was not seen in any of our test media. Instead, as seen in Figure 4, the LB data, the equivalent or faster growth conferred by pMB2 was not seen with the mini-plasmid pRMKO, which grew at a rate that was slower than equivalent-sized plasmid pLMJ50. Intriguingly, pRMC, the cloned deleted region also conferred slower growth than wildtype, although in this case the carrying cost can be attributed to the higher copy number of the clone. Attempts to move the large insert to a lower copy-number vector were not successful and therefore it was not possible to fully complement the mini-plasmid deletion in *trans*. We therefore elected to functionally characterize the *sitABCD* genes that are within the deleted sequence.

**Figure 4:**
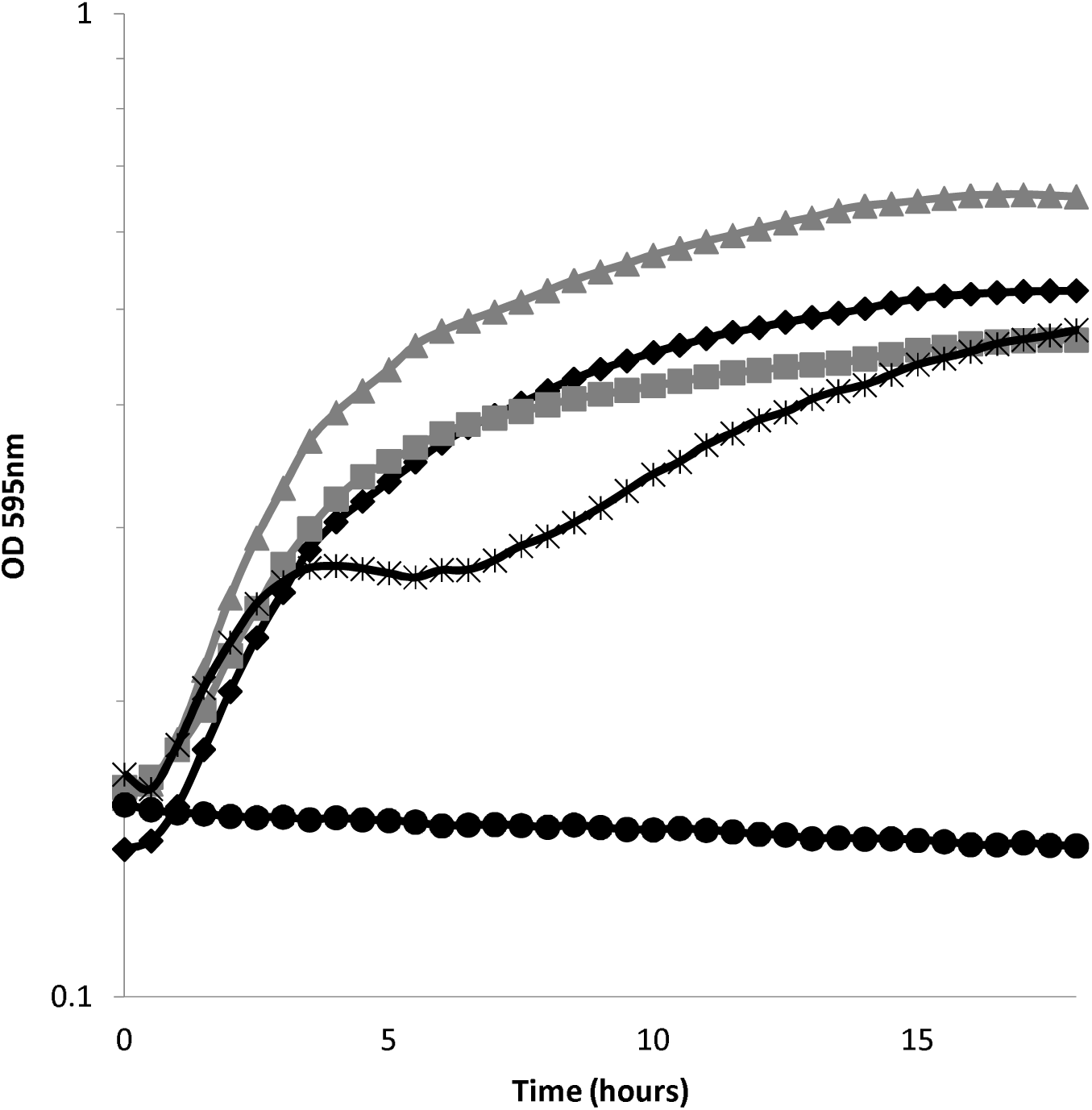
18 h growth time-course of DH5α carrying the pRMKO miniplasmid derived from pMB2 (black crosses) compared to growth of DH5α (black diamonds), DH5α carrying pMB2 (grey triangles) and 90 Kb control plasmid pLMJ50 (grey squares). Black circles represent a no bacterial control.

We amplified the *sitABCD* operon and its promoter from plasmid pSIT1 received from Laura Runyen-Janecky (46), using primers sitF and sitR and cloned them into pACYC177 to create pINK2301. In media that has been depleted by EDDA, DH5α grows much slower than M63c, the *E. coli* strain from which pMB2 was isolated. Transformation of DH5α with pMB2 produced a faster growth rate that was lower in DH5α (pRMKO). As shown in Figure 5, this growth defect could be complemented in *trans* with pINK2301.

**Figure 5:**
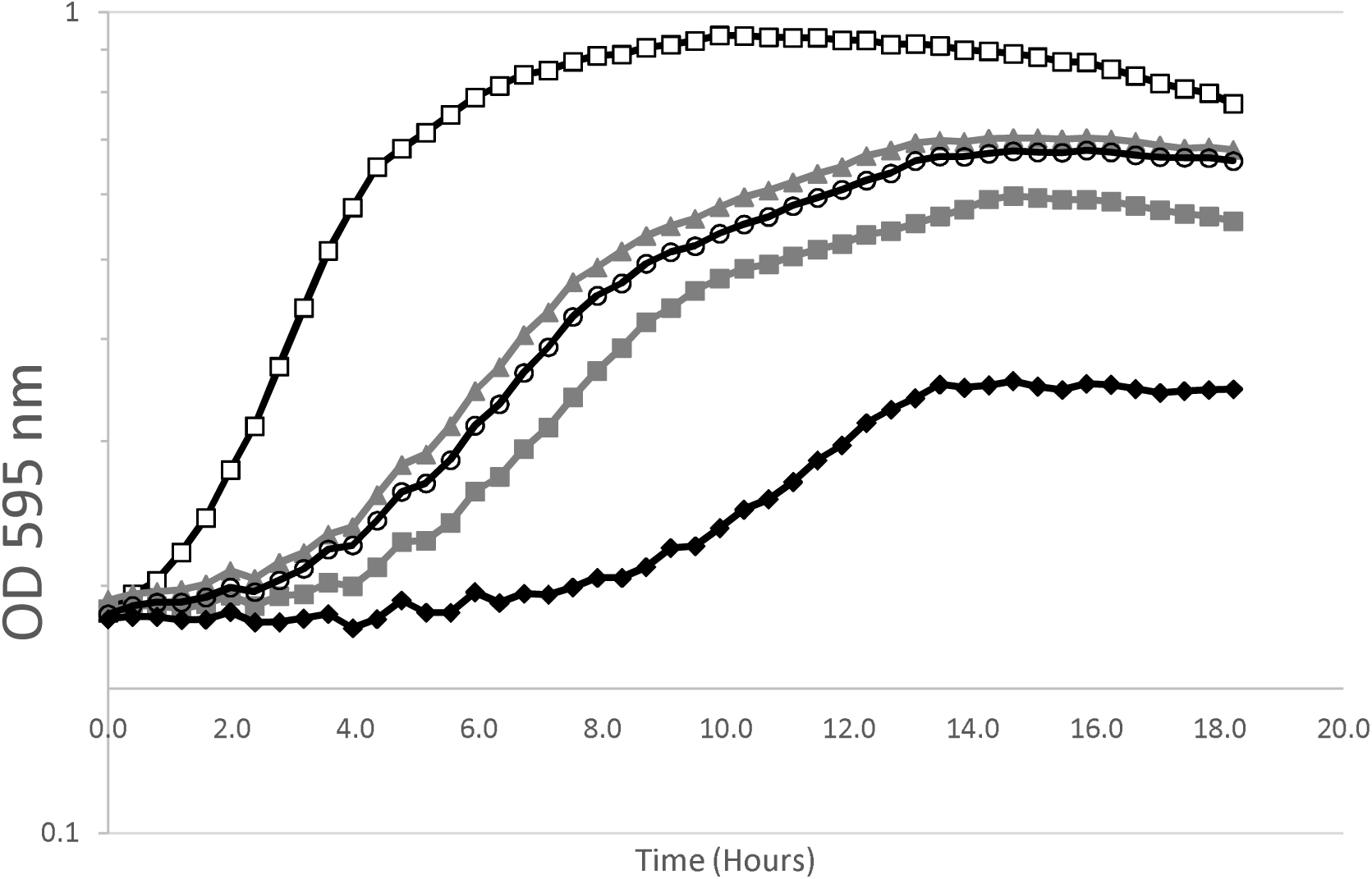
18 h growth time-course in David’s minimal media pre-treated with deferrated EDDA. Wildtype M63c strain (black open squares) compared to DH5α (black diamonds), DH5α carrying pMB2 (grey triangles), pRMKO miniplasmid (grey squares), pRMKO miniplasmid plus the cloned sit genes in pINK2301 and 90 Kb control plasmid pLMJ50 (grey squares).

To determine whether pMB2 enhanced bacterial fitness, we conducted competition experiments between strain M63c, the strain from which pMB2 was originally isolated, and commensal *E. coli* strain HS. We also competed DH5α strains carrying pMB2 and DH5α (pRMKO) against one another. These experiments were performed in rich media (LB) as well as in DMEM in the absence of antibiotics, however antibiotics were used to select competitors on plates to perform counts. In both media, strains carrying pMB2 outcompeted commensal or laboratory strains. Conversely, DH5α (pRMKO) suffered a competitive disadvantage against DH5α (pMB2), as well as against plasmidless DH5α (Figure 6).

**Figure 6:**
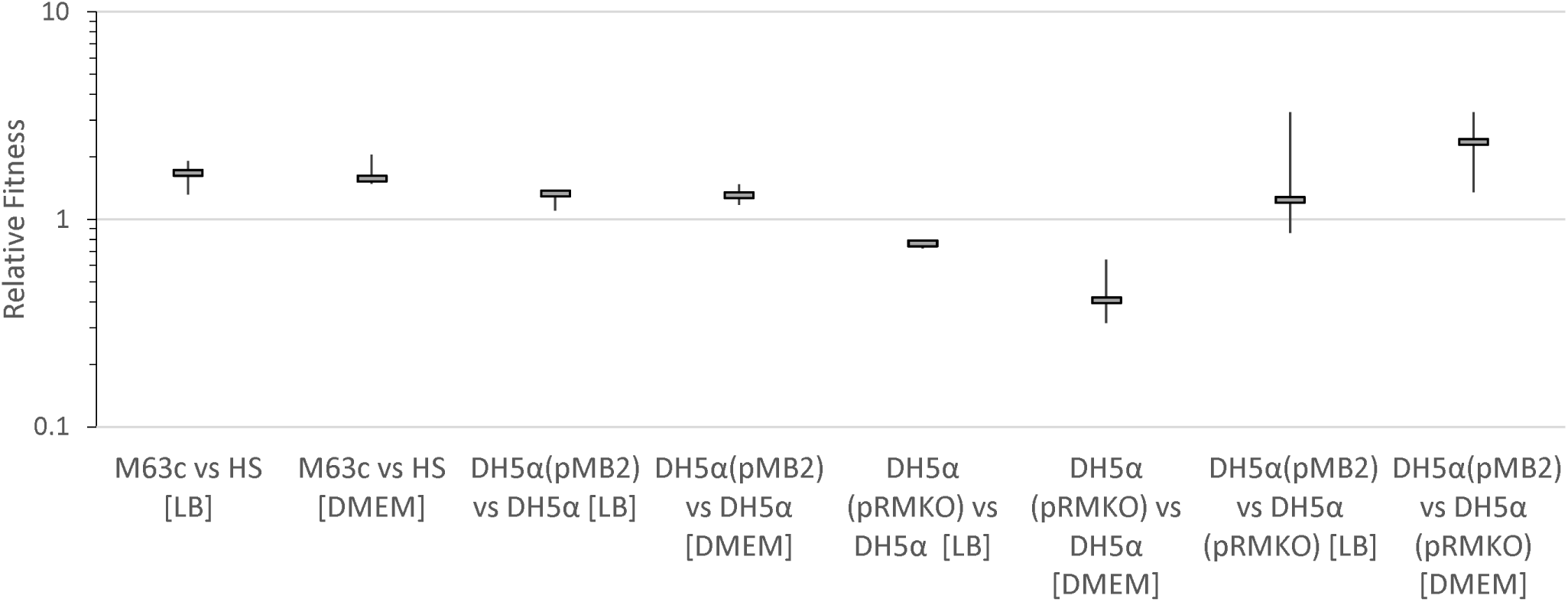
Relative fitness of the initially listed strain compared to the subsequently listed strain in rich media (LB) and minimal media (DMEM).

We sought to determine whether the plasmids were stably inherited and found that this was the case in source strain M63c over 200 generations. However, pMB2 and pRMKO were less stably inherited in a DH5α background, with pRMKO proving to be less stably inherited than pMB2 in the medium term (Figure 7).

**Figure 7:**
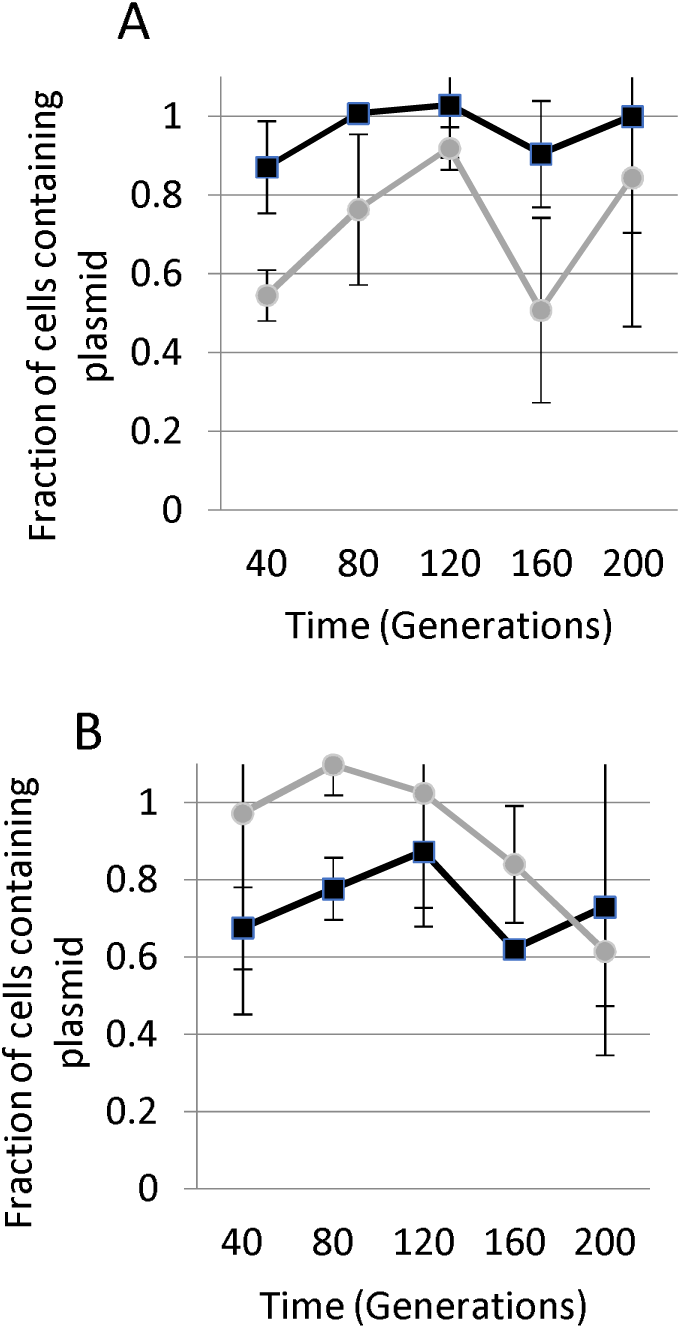
Plasmid stability in the absence of selection (a) stability of pMB2 in M63c, its natural host (dark line) and control plasmid pBR322 in DH5α (grey line) (b) stability of pMB2 (grey line) and pRMKO (dark line) in DH5α.

### A PefB homolog encoded on pMB2 interferes with autoaggregation

We observed that DH5α carrying the pRMKO mini-plasmid showed significant autoaggregation, particularly under low nutrient conditions. As shown in Figure 8a, autoaggregation was easily quantified in DH5α (pRMKO) but absent in DH5α (pMB2) cultures grown under the same conditions (p<0.001). Cloning the deleted region into pBluescript also reduced autoaggregation of DH5α compared to an isogenic strain carrying the vector alone (Figure 7a). We examined the sequence of the region deleted from pRMKO for candidate genes that could contribute to autoaggregation. Excluding hypothetical orfs, the best candidate is a *papB/pefB* homolog at position 11,258-11,530 of the plasmid. PapB and PefB are prototypical members of a family of mobile element-borne fimbrial regulators (61). They regulate core chromosomal adhesins in *Escherichia* (62, 63). We cloned the *pefB* homolog under the control of the arabinose promoter to produce pRMPefB. We then measured the ability of pRMPefB to complement the hyperautoaggregation phenotype produced by pRMKO. In the presence of arabinose, when the *pefB* homologue is induced, autoaggregation was diminished in DH5α (pRMKO, pRMPefB). This phenotype was not seen in the presence of glucose, which represses the arabinose promoter (Figure 8b). Based on this finding, we attribute the hyperautoaggregation phenotype conferred by pRMKO to deletion of the pMB2 *pefB*.

**Figure 8:**
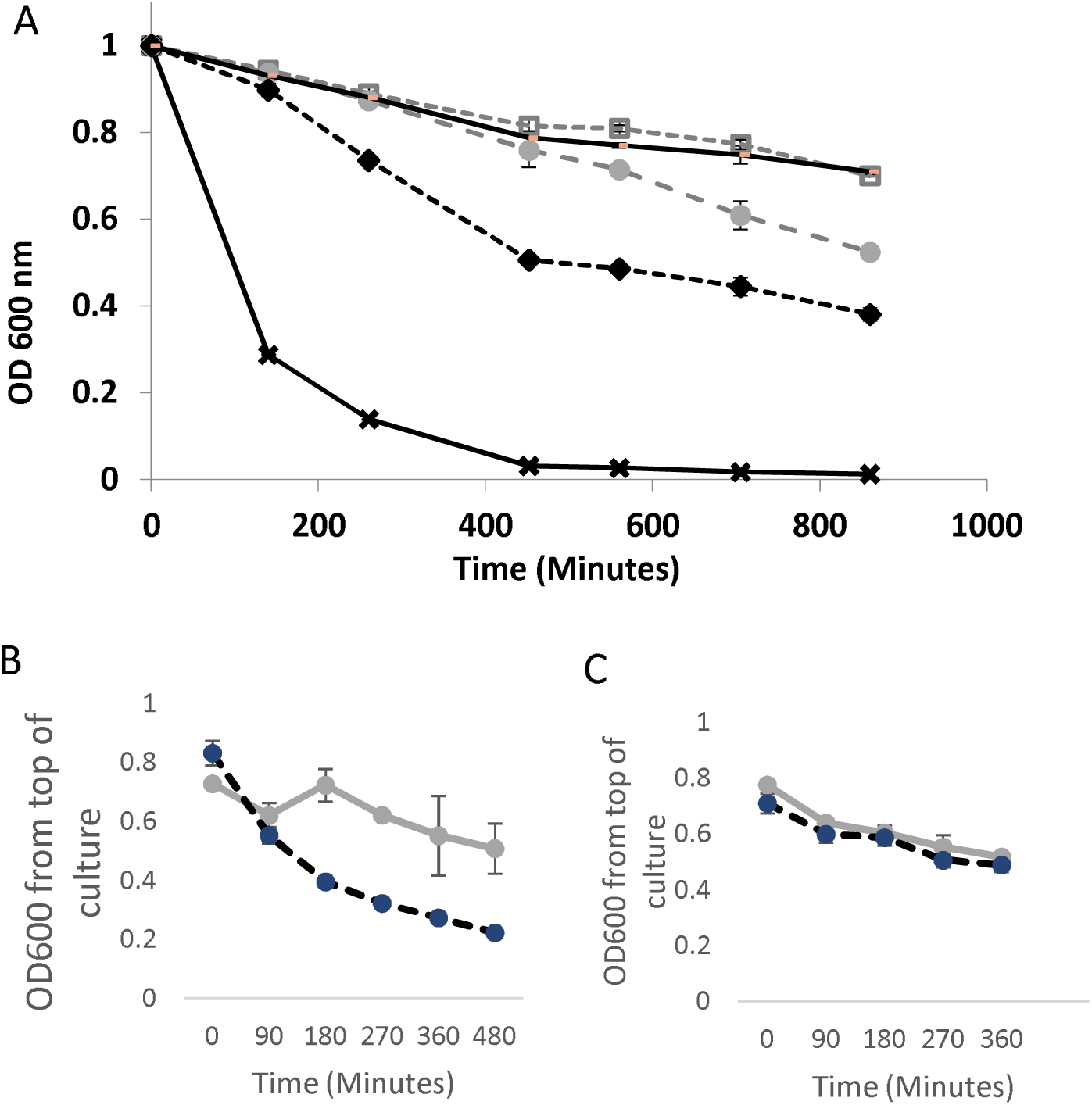
Autoaggregation in DMEM measured as absolute OD_600_ values from the top of a static culture sampled over time. (a) Autoaggregation of *E. coli* DH5α carrying pBluescript (grey closed circles on broken line), pMB2 (grey squares on broken line), pRMKO (black diamonds on broken line), pRMC (black line without marker) and pLMJ50 (black crosses). (b) *E. coli* DH5α carrying miniplasmid pRMKO and pINKpefB1, the *pefB* gene from pMB2 cloned under the control of the arabinose promoter. Autoaggregation was measured after growth in arabinose (solid grey line) or glucose (broken dark line) (b) *E. coli* ORN172 carrying pMB2 (solid grey line) and pRKMO (dark broken line).

PapB is capable of repressing the expression type 1 fimbriae and upregulating pyelonephritis-associated pili in uropathogenic *E. coli* (64). PefB is a plasmid-encoded PapB paralogue first detected in *Salmonella* Typhimurium plasmid that has similarly been shown to be able to repress type 1 fimbriae, albeit less efficiently (61). In DH5α (pMB2), PefB could be repressing chromosomally-encoded *fim* genes or other factors on pMB2 that are not deleted in pRMKO. Candidates for the latter include F (conjugative)-pili and, which are known to promote autoaggregation and biofilm formation (65). To determine which of these possibilities could be at play, we transformed pMB2 and pRMKO into strain ORN172, which is deleted for *E. coli fim* genes (66). In this Δ*fim* background, there were no *pefB*-associated differences in autoaggregation. Therefore interactions of *pefB* with chromosomal factors, and specifically pili is the most likely explanation for the hyperautoaggregation seen in DH5α (pRMKO). A repressor of *fimA* could enhance F-pilus-mediated conjugation. We tested this hypothesis by comparing conjugative transfer rates to strain EC1502 from DH5α (pMB2) and DH5α (pRMKO). We recorded a transfer rate of 7.94 × 10^-6^ for DH5α (pMB2) in solid matings whereas conjugation from DH5α (pRMKO) could not be detected at the limits of the assay. The low conjugation rate overall and the impossibility of detecting rates below 1 × 10^-8^ made it impossible to perform a complementation experiment. However, altogether the available data support the idea that *pefB* repressed genes could interfere with conjugative transfer of pMB2.

## Conclusion

Antimicrobial use is the principal driving force behind the current epidemic of antimicrobial resistance. However, research has shown that, once evolved, multidrug resistance plasmids can be maintained in the absence of selection (67). There is a dearth of fitness defects reported for resistance plasmids in the literature and the reasons why some of them are evolutionarily successful are not well understood (58, 68). This study demonstrates that the *aac(6’)-Ib-cr* gene of strain M63c is carried on a large multidrug resistance plasmid pMB2, which also bears a number of genetic loci that support its evolutionary success irrespective of antibiotic selection. The 125 Kb plasmid does not have a carrying cost *in vitro* and a 32 Kb region of the plasmid accounts for selective success. Included in this evolutionarily significant 32 Kb region are the *sitABCD* genes, which promote growth under iron-limited conditions, along with as yet unidentified loci that promote growth in rich and minimal media. pMB2 is self-transmissible by an F-type conjugative system that may function more effectively due to repression of chromosomal genes by a *pefB* homolog located elsewhere on the plasmid. It is noteworthy that plasmids that are almost identical to pMB2 have been isolated from different parts of the world including Columbia, Italy and the UK. The *sit* and *pefB* genes studied in our research are highly conserved among most of them and therefore the fitness they confer is also likely to be common. Overall, these data showcase that fitness conferring mobile elements associated with resistance can co-evolve with host bacteria to compensate the costs of their carriage and propagate its transmission. Without a carrying cost large multidrug resistance plasmids could persist in commensal *E. coli* even in the absence of antimicrobial pressure. Future work on resistance plasmids and elements should focus more intently on fitness genes in addition to those that encode drug resistance.

## Acknowledgements

INO is an African Research Leader award supported by the UK Medical Research Council (MRC) and the UK Department for International Development (DFID) under the MRC/DFID Concordat agreement that is also part of the EDCTP2 programme supported by the European Union. This work was also supported by a Branco Weiss Fellowship from the Society in Science, Zürich, Switzerland and by US National Science Foundation ‘Research at Undergraduate Institutions’ award MCB 0948460 both to INO, and by undergraduate research studentships from Haverford College. BWO was the recipient of an International fellowship for Africa from American Society for Microbiology. We thank Drs David C Hooper, Alfredo G Torres and Laura Runyen-Janecky for strains. We are grateful to Erin Remaly, Rotimi Dada and Jennifer Hofmann for technical assistance and to Lawrence Wang for helpful comments.

