## Supplemental table 1 for "A large self-transmissible plasmid from Nigeria confers resistance to multiple antibacterials without a carrying cost"

**Supplementary table 1: Oligonucleotide primers used in this study**

| **Target gene** | **Primer** | **Primer Sequence (5’ to 3’)** | **Purpose; amplicon size** | **Reference** |
| --- | --- | --- | --- | --- |
| *aac(6’)-Ib* |  | TTGCGATGCTCTATGAGTGGCTA | *aac(6’)-Ib* (all alleles) detection; 482 bp | (1) |
|  |  | CTCGAATGCCTGGCGTGTTT |  |  |
| *oqxA* |  | CTCGGCGCGGATGATGCT | *oqxA* detection; 392 bp |  |
|  |  | CCACTCTTCACGGGAGACGA |  |  |
| *qnrA* | qnrA-1A | TTC AGC AAG ATT TCT CA | *qnrA* detection; 608 bp | (2) |
|  | qnrA-1B | GGC AGC ACT ATT ACT CCC AA |  |  |
| *qnrB* | qnrB-CS-1A | CCT GAG CGG CAC TGA ATT TAT | *qnrB* detection, 389 bp | (2) |
|  | qnrB-CS-1B | GTT TGC TGC TCG CCA GTC GA |  |  |
| *qnrS* | qnrS-1A | CAA TCA TAC ATA TCG GCA CC | *qnrS* detection; 621 bp | (2) |
|  | qnr-1B | TCA GGA TAA ACA ACA ATA CCC |  |  |
| *qepA* | qepA-F | GCAGGTC CAGCAGCGGGTAG | *qepA* detection; 218 bp | (3) |
|  | qepA-R | CTTCCTGCCCGAGTATC GTG |  |  |
| *gyrA* | gyrA12004 | TGC CAG ATG TCC GAG AT | *gyrA* QRDR amplification | (4) |
|  | gyrA11753 | GTA TAA CGC ATT GCC GC |  |  |
| *parC* | EC-PAR-A | CTG AAT GCC AGC GCC AAA TT | *parC* QRDR amplification | (5) |
|  | EC-PAR-B | GCG AAC GAT TTC GGA TCG TC |  |  |
| *adk* | *adk*F | ATTCTGCTTGGCGCTCCGGG | MLST | (6) |
|  | *adk*R | CCGTCAACTTTCGCGTATTT |  |  |
| *fumC* | *fumC*F | TCACAGGTCGCCAGCGCTTC | MLST | (6) |
|  | *fumC*R | GTACGCAGCGAAAAAGATTC |  |  |
| *gyrB* | *gyrB*F | TCGGCGACACGGATGACGGC | MLST | (6) |
|  | *gyrB*R | ATCAGGCCTTCACGCGCATC |  |  |
| *icd* | *icd*F | ATGGAAAGTAAAGTAGTTGTTCCGGCACA | MLST | (6) |
|  | *icd*R | GGACGCAGCAGGATCTGTT |  |  |
| *mdh* | *mdh*F | ATGAAAGTCGCAGTCCTCGGCGCTGCTGGCGG | MLST | (6) |
|  | *mdh*R | TTAACGAACTCCTGCCCCAGAGCGATATCTTTCTT |  |  |
| *purA* | *purA*F | CGCGCTGATGAAAGAGATGA | MLST | (6) |
|  | *purA*R | CATACGGTAAGCCACGCAGA |  |  |
| *recA* | *recA*F | CGCATTCGCTTTACCCTGACC | MLST | (6) |
|  | *recA*R | TCGTCGAAATCTACGGACCGGA |  |  |
| *pefB* | *pefBF* | CACCACTCCCTCCCCCTATCCAA | Cloning the *pefB* orf; 273 bp | This study |
|  | *pefBR* | CTCACGGTAGAAATATTTAAGAGC |  |  |
| *sitABC* | *sitF* | CACCCTCTCAATAAAAAAAGTAACCATG | Cloning *sit* operon | This study |
|  | *sitR* | CTAATCGGATGATAACAAATCCC | 3.4 Kb |  |
| *16S rRNA* | *10F* | AGTTTGATCATGGCTCACATTG | Bacterial identification; 1.5 Kb | (7) |
|  | *1507R* | TACCTTGTTACGACTTCACCCCAG |  |  |

MLST – multi-locus sequence typing; QRDR – quinolone-resistance determining region
